# An in vivo model of Amyloid-Related Imaging Abnormalities (ARIA) predicts ARIA incidence of anti-Aβ antibodies seen in clinical trials

**DOI:** 10.64898/2026.01.30.702402

**Authors:** Edward D. Plowey, Jordan Abettan, Peter Cheng Te Chou, Ming Lye-Barthel, Ana Claudia Silva Amaral, Jennifer Sebalusky, Stefan Hamann, Jared Srinivasan, Roland Brown, Fang Qian, Paul Weinreb, Dominic M. Walsh, Daniel Bradley, Thierry Bussiere

**Affiliations:** Biogen, Cambridge, MA, USA

## Abstract

Amyloid-related imaging abnormality (ARIA) is the most common adverse event associated with amyloid-beta (Aβ) immunotherapy. The neuropathology underlying active ARIA lesions has been studied in only a few rare fatal cases. A faithful translational model of ARIA should allow a better understanding of pathogenic mechanisms and enable development of mitigation strategies. Since bapineuzumab induced the most frequent and severe forms of ARIA in humans, we assessed the ability of its murine precursor, 3D6, to induce ARIA-like lesions by applying clinically-validated MRI sequences in a mouse model with extensive amyloid deposition. Using this paradigm we documented ARIA-like changes based on their imaging features, location, and temporal evolution and established postmortem histopathologic correlates. We observed that ARIA-like lesions were ubiquitously induced with 3D6, including T2-hyperintense leptomeningeal effusions, T2-hyperintense parenchymal lesions and T2* hypointense parenchymal lesions. Histological assays demonstrated meningovascular inflammation as well as vascular mural permeation and microvascular lesions, including microhemorrhages and microinfarcts. Compared to 3D6-treated mice, animals treated with mouse analogues of aducanumab and gantenerumab showed less frequent radiologic and histopathologic ARIA-like lesions with delayed onset. Our results align well with the differential incidence of ARIA observed in humans treated with different anti-Aβ antibodies. Collectively, our studies demonstrate that: i) anti-Aβ antibodies chronically administered to 5xFAD mice can induce MRI lesions reminiscent of human ARIA; ii) this paradigm allows identification of transient lesions other than microhemorrhages, and (iii) the incidence and severity of ARIA-like lesions induced with three clinically tested antibodies faithfully recapitulates that seen in humans.

**One Sentence Summary:** MRI and histological correlation in a preclinical ARIA mouse model treated with anti-Aβ antibodies that recapitulates results seen in humans.

## INTRODUCTION

Treatment of Alzheimer’s disease (AD) patients with certain anti-Aβ antibodies is known to cause magnetic resonance imaging (MRI) signal abnormalities referred to as amyloid-related imaging abnormalities (ARIA) (*1, 2*). ARIA detected as T2-FLAIR hyperintensities indicates the presence of edema or sulcal effusions resulting from abnormal water and/or protein content in the brain parenchyma or subarachnoid space and are referred to as ARIA-edema/effusion (ARIA-E). ARIA detected as T2* (gradient echo, GRE) hypointensities indicates the presence of microhemorrhages in the brain parenchyma or superficial siderosis in the subarachnoid space and are referred to as ARIA-microhemorrhages/superficial siderosis (ARIA-H). Case reports of human postmortem brain examinations in rare fatal cases of ARIA have identified underlying Aβ-related angiitis (*3-5*), suggesting a mechanistic link between cerebral amyloid angiopathy (CAA), immunotherapy-induced CAA inflammation and ARIA. Since tissue from subjects with active ARIA events is not widely available, the generalizability of pathologic findings in rare fatal ARIA cases to the common spectrum of asymptomatic and symptomatic ARIA is unclear. In the most recent report of two subjects with aducanumab-induced ARIA, brain examination was performed years after the active ARIA events (*6*).

Preclinical models may be useful to identify histopathologic changes and pathogenic mechanisms underlying ARIA. Neuropathologic studies of Amyloid Precursor Protein (APP) transgenic mouse models chronically treated with Aβ immunotherapy have previously reported inflammation of meningeal blood vessels (meningovascular inflammation, MVI) and microhemorrhages (*7-14*). MVI has been hypothesized to cause ARIA-E via the breakdown of vascular mural integrity and subsequent micropermeation of brain parenchyma by proteinaceous fluids and contents, while the detection of hemosiderin is considered to be a correlate to microhemorrhage found in ARIA-H. However, with the notable exception of Luo and colleagues who focused on correlation of microhemorrhages detected in ex vivo T2* sequences and histology in 17-23-month-old Tg2576 mice (*15*), ARIA-neuropathologic correlations in APP transgenic mouse models have not been reported.

The current manuscript reports a series of studies in which 9.4T MRI scans were performed longitudinally to detect ARIA-like events in 5xFAD mice (n=146; Table 1) chronically treated with anti-Aβ monoclonal antibodies (mAbs). We describe the MRI characteristics of ARIA-like lesions in these mice, many of which showed similarities to ARIA-E and ARIA-H lesions that occur in patients. Furthermore, we correlate brain histopathologic changes, including MVI, vascular mural permeation and microhemorrhage/hemosiderin, with MRI events. Our findings lend insights into the neuropathologic mechanisms underlying ARIA in this model and identify pathologic endpoints relevant for the development of ARIA mitigation strategies for anti-Aβ immunotherapeutics.

**Table 1.** Summary of experimental treatment studies with Aβ antibodies. A total of 153 mice (146 5xFAD and 7 WT) were used in 7 separate experiments. Numbers in parentheses are milligrams per kilogram for weekly intraperitoneal doses. *includes 7 WT mice that were treated with 3D6 and showed no ARIA and no vascular inflammation.

| Studies | Age (months) | Controls |  | 3D6 (mg/kg) |  |  |  | Aducanumab (mg/kg) |  | Gantenerumab (mg/kg) |  |  |  |  |
| --- | --- | --- | --- | --- | --- | --- | --- | --- | --- | --- | --- | --- | --- | --- |
|  |  | Untreated | Isotype | (5) | (10) | Agly(10) | <sup>ch</sup>(5) | <sup>ch</sup>(50) | (50) | (5) | Agly(5) | (10) | (30) | (50) |
| 1 | 8 or 14 | - | - | 19* | - | - | - | - | - | - | - | - | - | - |
| 2 | 7-8 | 10 | 8 | - | 8 | 8 | - | - | - | - | - | - | - | - |
| 3 | 13 | - | - | 5 | - | - | - | - | - | 5 | 5 | - | - | - |
| 4 | 11 or 15 | - | - | 6 | - | - | - | - | - | 6 | - | 6 | 6 | - |
| 5 | 8 | - | - | 5 | - | - | - | - | - | - | - | - | - | 10 |
| 6 | 7, 9 or 13 | - | - | 4 | - | - | - | - | - | - | - | - | 7 | 5 |
| 7 | 9.5 | - | - | 6 | - | - | 6 | 9 | 9 | - | - | - | - | - |

## RESULTS

### ARIA-like lesions detected in 5xFAD mice treated with 3D6 by MRI sequences used to monitor ARIA in humans

The characteristics of ARIA seen in clinical trials with different anti-Aβ mAbs is distinguishable, only by the incidence and severity of the lesions. Treatment of patients with bapineuzumab induced the most frequent occurrences and severe forms of ARIA (*16-18*). Thus, we assessed the ability of the murine precursor of bapineuzumab (3D6) to induce ARIA-like lesions using MRI sequences validated in humans and applied to a mouse model with extensive parenchymal and vascular amyloid deposition (*19-21*). T2w and T2* MRI sequences used for monitoring ARIA in patients treated with anti-Aβ immunotherapy were performed at baseline and then weekly upon chronic treatment of 5xFAD mice with 3D6. ARIA-like lesions are defined based on their imaging features, location, and temporal evolution.

ARIA-like lesions appear as hyperintense or hypointense signals using a T2w MRI sequence. T2w-hyperintense lesions are either focal, appearing as bright and well-circumscribed, or diffuse and often occupying a large brain volume (**Figs 1A-B**). Focal T2w-hyperintense parenchymal lesions sometimes emerged in regions where diffuse lesions were previously described (**Fig. 2**). T2w-hyperintense lesions are also detected in the leptomeningeal space and are defined as subarachnoid convexity lesions (**Fig. 1C**). Focal or diffuse parenchymal lesions are reminiscent of ARIA-E edema observed in patients, whereas convexity lesions are akin to sulcal effusions observed in patients. T2-hypointense focal lesions can be detected using T2w or T2* sequence. They are usually small in size and well-circumscribed within the brain parenchyma, and are less frequent than T2w-hyperintense lesions (**Fig. 1D**). In some cases, T2w-hyperintense lesions changed from week-to-week with many lesions being transient and resolving over the course of the study period, but others evolved into T2w/T2*-hypointense cortical lesions (**Fig. 2**). Multiple T2 lesions with any of the features described above may appear throughout the brain of the same animal during chronic treatment (**Fig. 2**), or they may coexist at a given timepoint (**Fig. 3**).

**Figure 1.**
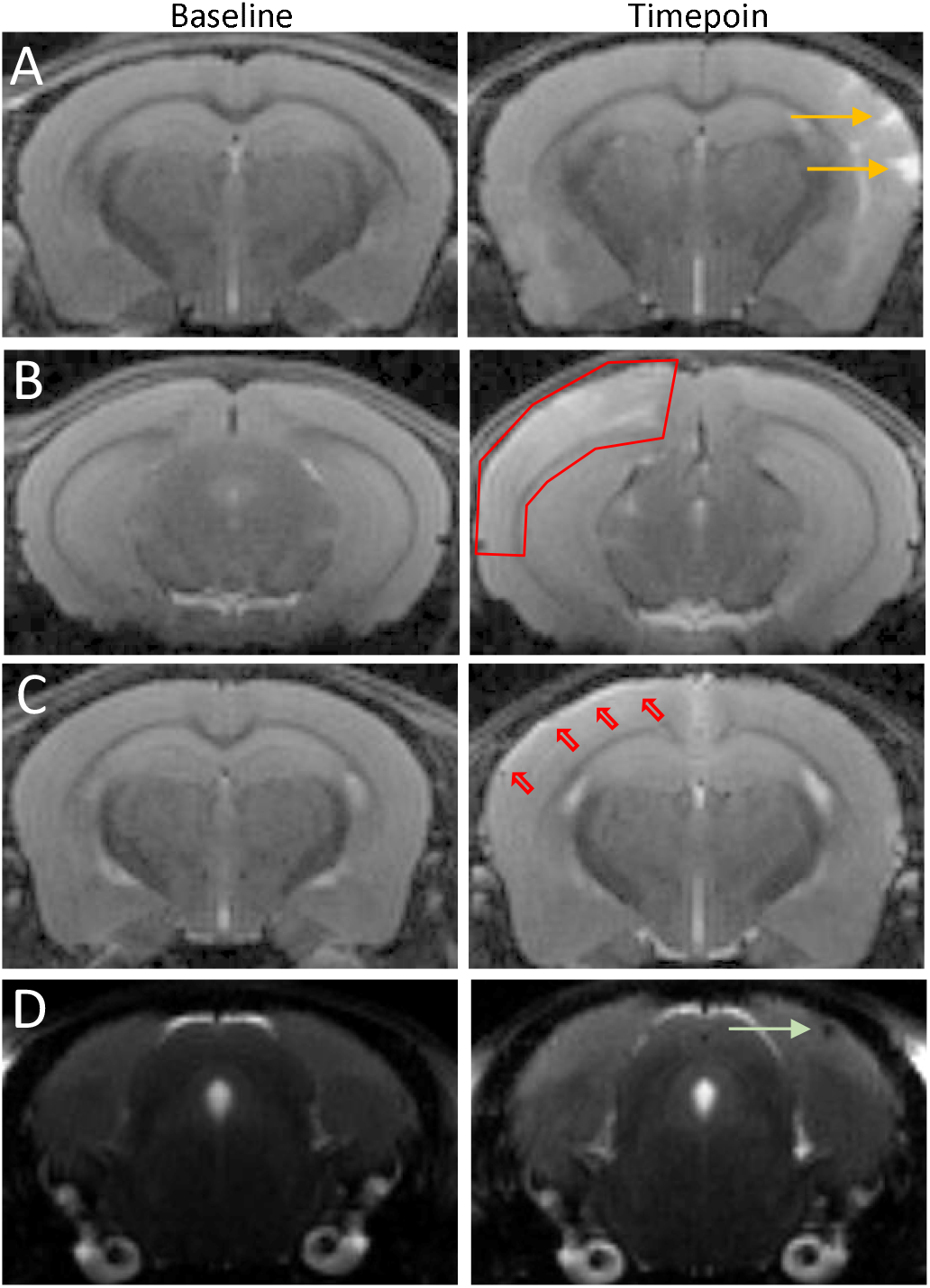
ARIA-like lesions observed in 5xFAD mice treated with 3D6 mIgG2a using T2w MRI sequence. Representative T2-weighted MRI scans depicting different types of ARIA-like lesions observed in animals treated with weekly doses of 3D6 mIgG2a (5 mg/kg; Study 1), are shown. Absence of ARIA-like lesions was verified prior to initiation of weekly dosing and imaging (Baseline, left column) (**A).** T2w-hyperintense focal and well-circumscribed cortical lesions, left side (orange arrows) after 6 weekly doses (Animal 16, week 6, Slice 14). (**B)**. T2w-hyperintense diffuse cortical lesion occupying a large brain volume, right side (red frame) after 6 weekly doses (Animal 11, week 6, Slice 11). (**C)**. T2w-hyperintense subarachnoid convexity lesion (red arrows) observed on the cortical surface, right side, after 3 weekly doses (Animal 22, week 3, Slice 14). (**D)**. Small T2w-hypointense focal cortical lesion, left side (green arrow) observed after 5 weekly doses (Animal 4, week 5, Slice 8). 30 MRI slices (0.5 mm) were acquired for each animal and were numbered in increasing order in the caudo-rostral direction.

**Figure 2.**
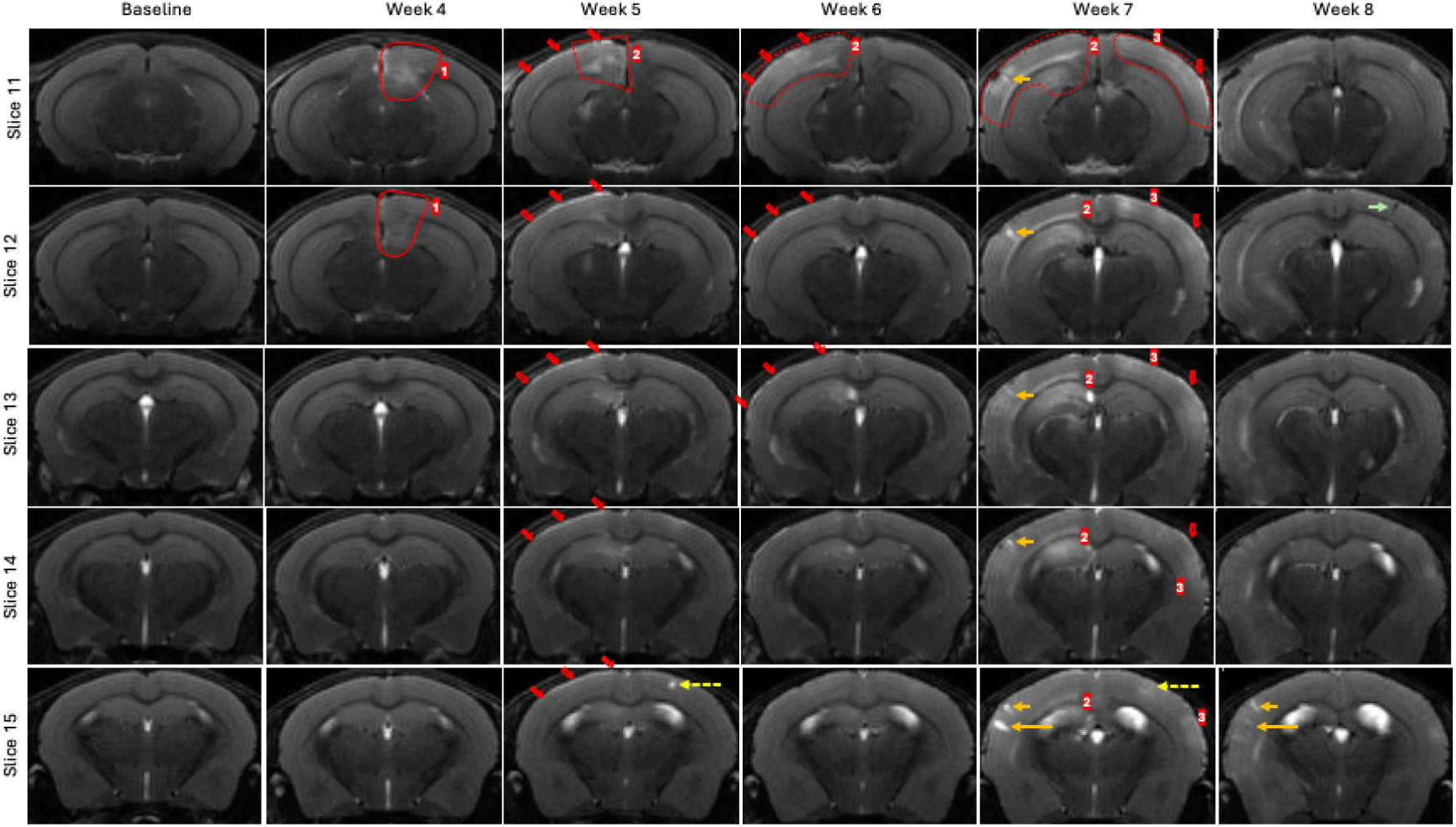
Temporal evolution of ARIA-like lesions in a 5xFAD mouse treated with 3D6 mIgG2a using T2w MRI sequence. T2-weighted MRI baseline and weekly scans between week 4 and end of study (week 8) showing evolution of multiple ARIA-like lesions observed in animal 11/study 1 treated with weekly doses of 3D6 mIgG2a (5 mg/kg). Absence of ARIA-like lesions at baseline was verified at all levels (left column). ARIA-like lesions are reported in slices 11 to 15 to illustrate possible evolution of lesions over time and might not represent the entirety of the volume of the lesions. Several occurrences of T2w-hyperintense diffuse ARIA-like lesions with different types of evolution were identified through the last 5 weeks of the study: ARIA-like lesion 1 (red frame 1) was first observed after 4 weekly doses (week 4) in the left retrosplenial region (Slices 10-11-12) and had resolved within one week (week 5). ARIA-like lesion 2 (red frame 2) first appeared in the right retrosplenial region after 5 weekly doses (week 5) and then extends further into the neocortex and hippocampal formation as treatment continues in the following weeks. The intensity of the diffuse signal seemed to decrease by the end of the study (week 8), but noticeably several focal hyperintense lesions appeared within the same area (orange arrows; week 7). These latter lesions evolve rapidly, with most of them resolving within a week. ARIA-like lesion 3 (red frame 3) appeared as a diffuse hyperintense signal involving a large area of the left neocortex after 7 weekly doses (week 7; slice 11). Most of the signal resolved within a week, but few isolated areas of brighter signal remain (week 8; slices 12-15). A single focal T2w-hyperintense signal detected in the left th cortex after the 5 dose (dashed yellow arrow, week 5; slice 15) resolved rapidly and only a weak outline of the signal remains visible 2 weeks later (week 7; slice 15). A small focal T2w-hypointense ARIA-like lesion is detected in the left cortex in a region previously involved with a bright diffuse signal (green arrow, week 8; slice 12), illustrating the potential evolution of T2w edema type of lesion into a microhemorrhage-type of lesion. A right side T2w-hyperintense subarachnoid convexity lesion (red arrows) is mostly visible after the 5 ^th^ dose (week 5, slices 11-15) and had resolved by the end of the study A similar type of ARIA-like lesion with a lower intensity appeared in the left hemisphere after the 7 ^th^ dose (red arrow, week 7; slices 11-14) and seemed to have resolved by the end of the study.

**Figure 3.**
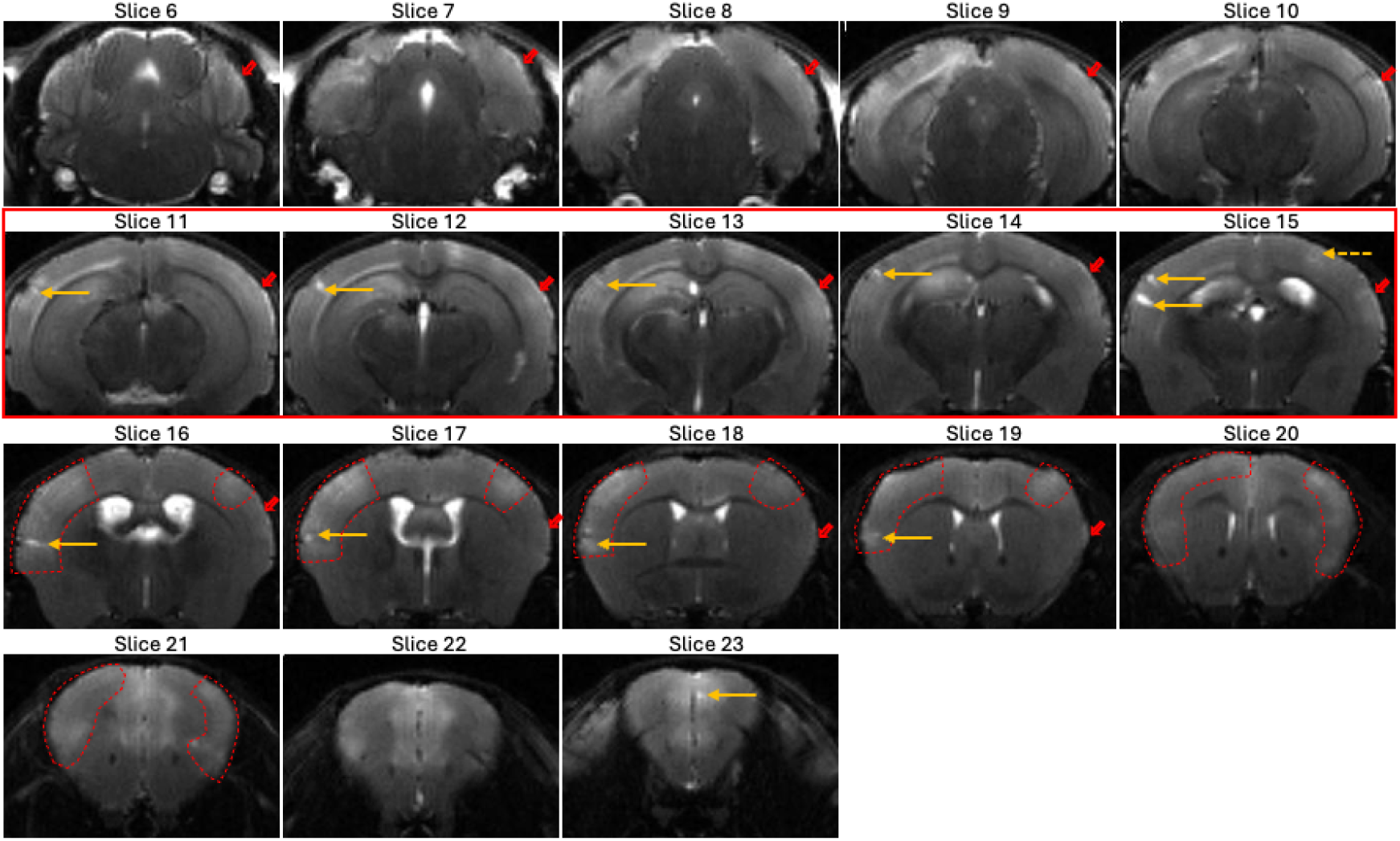
Anatomical distribution of ARIA-like lesions in a 5xFAD mouse treated with 3D6 mIgG2a using T2w MRI sequence. T2-weighted MRI scans spanning slices 6 to 23 at week 7 showing ARIA-like lesions observed in animal 11/study 1 treated with weekly doses of 3D6 mIgG2a (5 mg/kg). Different types of T2w ARIA-like lesions are randomly distributed through the entire brain, namely subarachnoid convexity lesion (red arrow), focal cortical lesions (orange arrows), diffuse cortical lesions (dashed red shapes; slices 16-21). Most of the T2w-hyperintense lesions are visible in more than one MRI scans. Scans presented in second row (slices 11-15; red frame) are the same than in figure 2.

ARIA-like lesions were very rarely observed in untreated 5xFAD mice at baseline, and were not induced by treatment with isotype control murine IgG2a in 5xFAD mice or in wild-type mice treated with 3D6. ARIA-like lesions were most commonly observed after the 3^rd^ to 5^th^ dose, and ubiquitously occurred after the 10^th^ dose for an overall incidence of ∼100% in 5xFAD mice treated with 3D6. Lesions appear earlier when older mice with higher amyloid burden were treated with 3D6, whereas the prevalence of ARIA-like lesions in study 1 was lower in 8-month-old 5xFAD mice (3/7 mice) compared to 14-month-old 5xFAD mice treated with 3D6 (7/7 mice). Similarly ARIA-like lesions appear earlier (**Fig S1**) but are also more severe histologically if the 3D6 dose is increased from 5 to 10 mg/kg. These 3D6-induced severe lesions share some of the pathological characteristics of the severe cases of human ARIA, such as prominence of vascular inflammation.

### Meningovascular inflammation is the predominant underlying pathology for ARIA-like lesions in 5xFAD mice treated with anti-Aβ antibodies

In 5xFAD mice treated with 3D6 mIgG2a, histopathologic sections corresponding to sulcal and diffuse cortical T2w-hyperintense ARIA-like lesions usually showed no definitive changes in cortical parenchyma but frequently showed conspicuous multifocal meningovascular inflammation (MVI) lesions involving the cerebral convexities (**Fig. 4A B**). We employed a grading scheme (see Methods) to describe the nature and severity of MVI lesions: uncommonly, we observed perivascular clusters of mononuclear cells with no clear macrophage differentiation (Grade 1; **Fig. 4C**). Most often, MVI was comprised of macrophages with elongate or epithelioid morphology, the latter of which occasionally demonstrated morphologic evidence consistent with phagocytosed amyloid (**Fig. 4D, E**) and which was assigned a grade of 2 if the lesion was visible only at high power and grade 3 if the lesion was visible at low power. We frequently encountered fibrinoid change, indicative of endothelial injury, in inflamed vessels (**Fig. 4F**) which was assigned a grade of 4. Grade 4 was also assigned in rare cases in which neutrophils were observed among severe MVI infiltrates with fibrinoid change (**Fig. S2**), particularly in experiments in which animals whose ARIA onset was close to the time of the end of the dosing regimen and necropsy.

**Figure 4.**
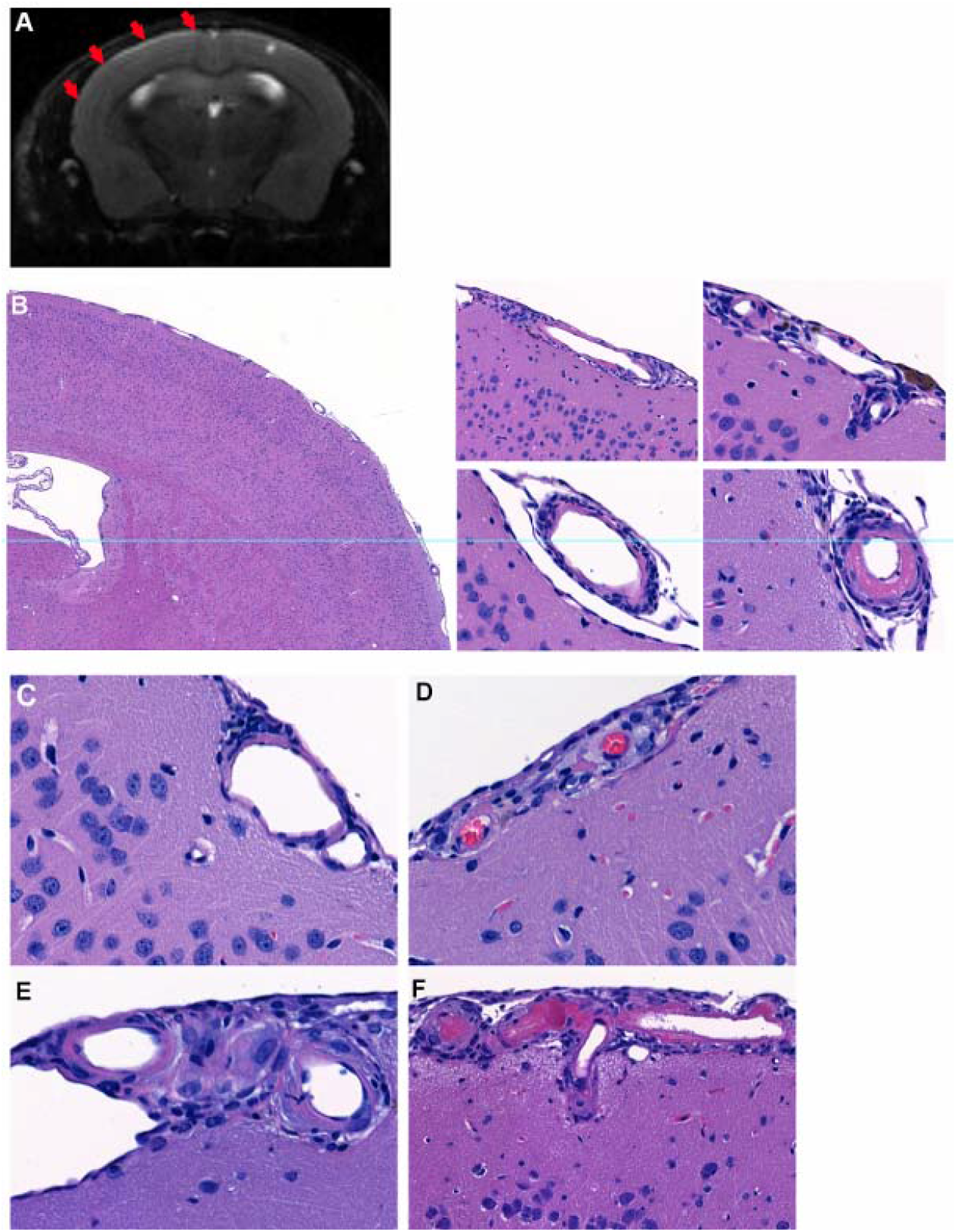
3D6 immunotherapy induces T2w-hyperintense cortical ARIA-like lesions that are associated with meningovascular inflammation. (**A).** T2-weighted MRI scan (Slice 15) from Animal 11/study 1, treated with weekly doses of 3D6 mIgG2a (5 mg/kg), showing at week 5: subarachnoid T2-bright signal consistent with convexity effusions (red arrows), right hemisphere, with sparse superficial cortical involvement by diffuse T2w-brightness; focal T2w-bright lesion in the superficial left cortex. (**B)**. H&E-stained sections of frontal cortex from animal 11 demonstrate meningovascular inflammation of varying severity involving many cortical arterioles. Definitive cortical parenchymal lesions were not identified in these sections. Left panel (original magnification 20x); right panels are 4 high power views (original magnifications 400x) of cortical arteriolar inflammation. (**C-F):** examples of MVI grading scheme employed in this study. A perivascular presence of mononuclear cells without obvious phagocytic morphology was designated as grade 1, minimal MVI. (**D):** Vascular macrophages that were clearly visible only at high power was designated as grade 2, mild MVI. (**E):** Multiple layers of vascular macrophages that were evident at low-to-medium power was designated as grade 3, moderate MVI. (**F)**. Vasculitis with fibrinoid change was designated as grade 4, severe MVI.

Over all experiments, compared to mice with Grades 0-1 MVI which showed a probability of ARIA of 0.018 (95% CI 0.002 to 0.128), odds of ARIA-E in mice with Grades 2, 3, and 4 MVI were 44 (95% CI 2.68 to 722), 342 (95% CI 13.3 to 8810) and 3430 (95% CI 35.2 to 334,000) times higher, respectively (**Table 2**). An odds ratio of 78 (95% CI 1.97 to 3090) was observed for mice with Grade 4 MVI compared to mice with Grade 2 MVI. These results substantiate a significant role for MVI presence and severity as a key pathologic determinant of ARIA in 5xFAD mice treated with Aβ antibodies. Moreover, sensitivity and specificity analysis demonstrated that histologic detection of any grade of MVI was highly sensitive (99%) and moderately specific (74%) for ARIA identified on MRI (**Table 3**). Stacked bar charts of MVI grade and ARIA by experiment (**Fig. 5**) illustrate the strong association between MVI grade and the presence of ARIA at the study level, with several instances of low-grade MVI occurring without ARIA suggesting that MVI emerges first. Notably, several animals in the gantenerumab and aducanumab groups demonstrated minimal to mild, and occasionally focally moderate, MVI without ARIA detection.

**Table 2.**
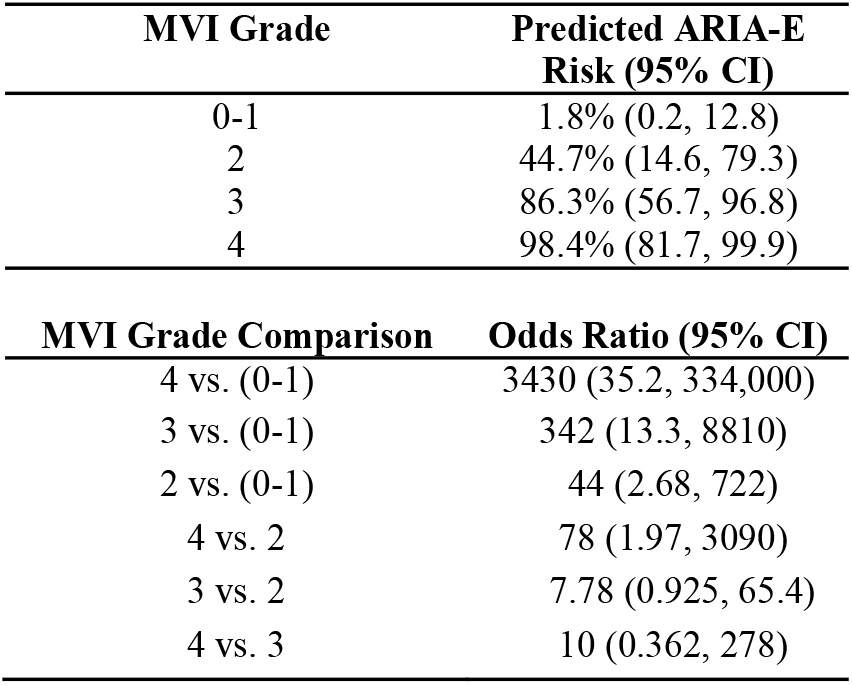
Mixed effects logistic regression model results: predicted ARIA-E risk by MVI Grade, and pairwise comparisons between MVI grades.

| MVI Grade | Predicted ARIA-E Risk (95% CI) |
| --- | --- |
| 0-1 | 1.8% (0.2, 12.8) |
| 2 | 44.7% (14.6, 79.3) |
| 3 | 86.3% (56.7, 96.8) |
| 4 | 98.4% (81.7, 99.9) |

| MVI Grade Comparison | Odds Ratio (95% CI) |
| --- | --- |
| 4 vs. (0-1) | 3430 (35.2, 334,000) |
| 3 vs. (0-1) | 342 (13.3, 8810) |
| 2 vs. (0-1) | 44 (2.68, 722) |
| 4 vs. 2 | 78 (1.97, 3090) |
| 3 vs. 2 | 7.78 (0.925, 65.4) |
| 4 vs. 3 | 10 (0.362, 278) |

**Table 3.** Sensitivity and specificity analysis of MVI grade in ARIA.

| MVI Grade | Sensitivity % | Specificity % |
| --- | --- | --- |
| Grade 4 | 40 | 100 |
| ≥ Grade 3 | 83 | 97 |
| ≥ Grade 2 | 99 | 82 |
| ≥ Grade 1 | 99 | 74 |

**Table 4.** Comparison of ARIA characteristics in human and in mouse model.

| ARIA | Human | Mouse Model |
| --- | --- | --- |
| **Imaging**<br>• ARIA-E<br>• ARIA-H | • FLAIR<br>• GRE-T2* | • T2w / DWI<br>• T2* |
| **Pathology**<br>• ARIA-E<br>• ARIA-H | • Leakage of proteinaceous fluid<br>• Leakage of heme products | • Leakage of Albumin and mIgG<br>• Leakage of heme products (hemosiderin/Perls staining) |
| **Features** | • Different incidence with different mAbs (Bapineuzumab>>>Aducanumab~Gantenerumab)<br>• Dose-dependent<br>• ARIA-E transient, resolve spontaneously, or evolve to ARIA-H<br>• Isolated ARIA-H<br>• Recurrent (low frequency, different location) | • Different incidence with different mAbs (3D6>>><sup>ch</sup>Adu~<sup>ch</sup>Gant)<br>• Dose-dependent<br>• ARIA-E transient, resolve spontaneously, or evolve to ARIA-H<br>• Isolated ARIA-H<br>• Recurrent (different location) |
| **Comments** | • Probable role for CAA, vascular leakage<br>• Likely role for neuroinflammation:<br>• similarity with CAA-ri<br>• Reduced ARIA with BBB-shuttle anti-Aβ (Trontinemab)<sup>(22)</sup> | • Probable role for CAA, vascular leakage<br>• Likely role for neuroinflammation:<br>• No ARIA with antibodies with reduced effector function<br>• Meningovascular inflammation commonly observed<br>• Reduced ARIA with BBB-shuttle anti-Ab (ATV:Aβ)<sup>(20)</sup> |

**Figure 5.**
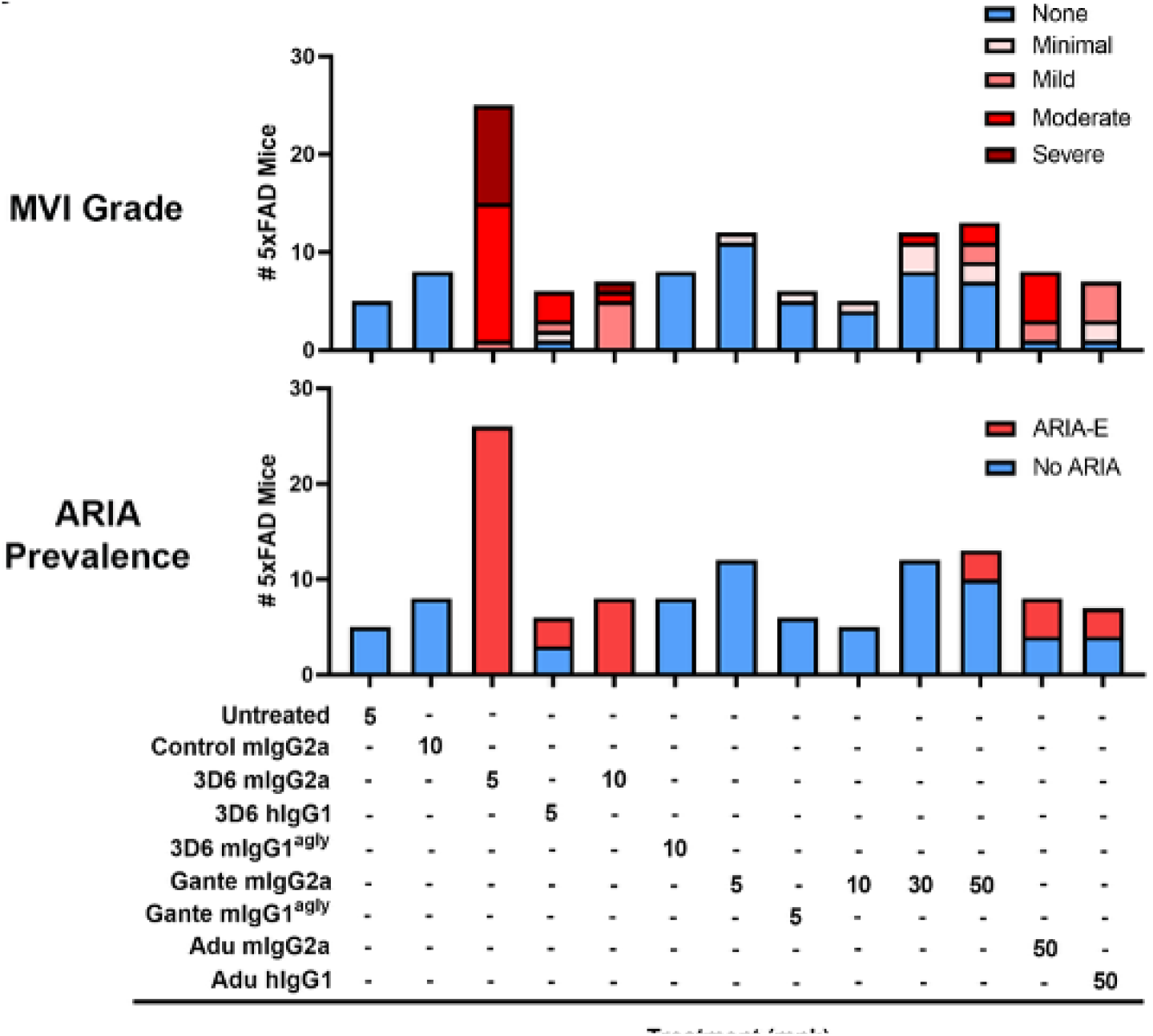
Occurrence of ARIA-like lesions is tightly associated with moderate to severe MVI. A strong association between prevalence of ARIA and prevalence of moderate-to-severe MVI was observed across treatment with the different antibodies. ARIA-like events were most frequent in 3D6-treated animals, where the most severe MVI were observed. Mild levels of MVI were found in some mice without presence of any detectable ARIA, most often in gantenerumab- and aducanumab-treated animals. MVI grade: each bar represents the total number of animal for the treatment groups listed below, with different color code allocated to each MVI grade; ARIA prevalence: each bar represents the total number of animals allocated to the treatment group listed below, with red sub-category indicating the number of animals with at least one ARIA-like event and blue sub-category the number of animals with no ARIA-E through the entire duration of the study.

Focal T2w-hyperintense parenchymal ARIA-like lesions could be observed to emerge in regions in which diffuse T2w-hyperintensities were previously detected and sometimes acute or subacute microinfarcts were observed by histopathology (**Fig. 6A, C, D**). These ARIA-like lesions often showed an elongate shape that was oriented perpendicularly to the cortical surface and sometimes demonstrated restricted diffusion (**Fig. 6B**). In addition to cortex, microinfarcts were sometimes observed in hippocampus, thalamus and cerebellar cortex. Microinfarct counts in all studies are plotted in **Fig. S3**.

**Figure 6.**
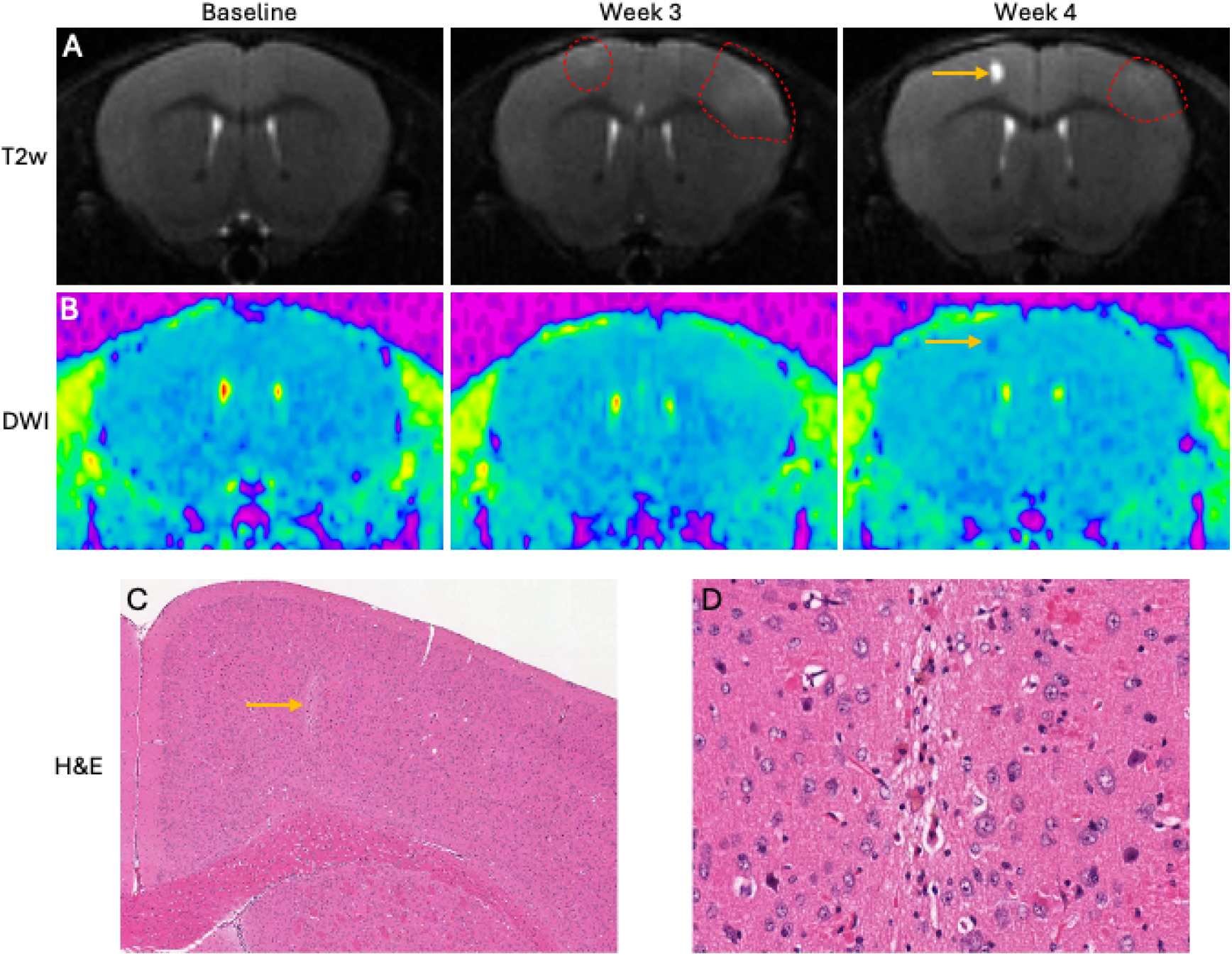
T2w-hyperintense, diffusion-restricted ARIA-like lesions observed in 5xFAD mice treated with 3D6 mIgG2a that correspond to microinfarcts on histopathology. **(A-B).** T2-weighted MRI scans (Slice 20) from Animal 21/study 2, treated with weekly doses of 3D6 mIgG2a (10 mg/kg), showing: emergence of bilateral diffuse T2w-bright cortical lesions (red shape) after 3 doses (week 3), persisting on the left side after the 4^th^ dose (week 4), or evolving into a well-circumscribed cortical T2w-bright lesion with restricted diffusion (DWI) on the right side (yellow arrow). (**C-D)**. H&E-stained sections from Animal 21 following the week 8 necropsy demonstrate a subacute cortical microinfarct corresponding to the T2-bright focal lesion with restricted diffusion (C, original magnification 50x; D, original magnification 400x).

T2*-hypointense cortical lesions were commonly observed following the resolution of poorly circumscribed diffuse T2w-hyperintense lesions (**Fig. 7A**). These ARIA-like lesions were either small with a punctate appearance, or linear and oriented perpendicularly to the cortical surface in a pattern consistent with perivascular hemosiderin surrounding penetrating cortical arterioles. Consistent with this impression, perivascular hemosiderin was observed on corresponding H&E-stained sections and Perls iron-stained sections (**Fig. 7B E**). Hemosiderin deposits were also commonly seen in the cortical molecular layer, closely subjacent to leptomeningeal arterioles, and were comprised of small scattered deposits consistent with mild blood leakage. A large acute hemorrhage was observed in only one animal (**Fig. S4**).

**Figure 7.**
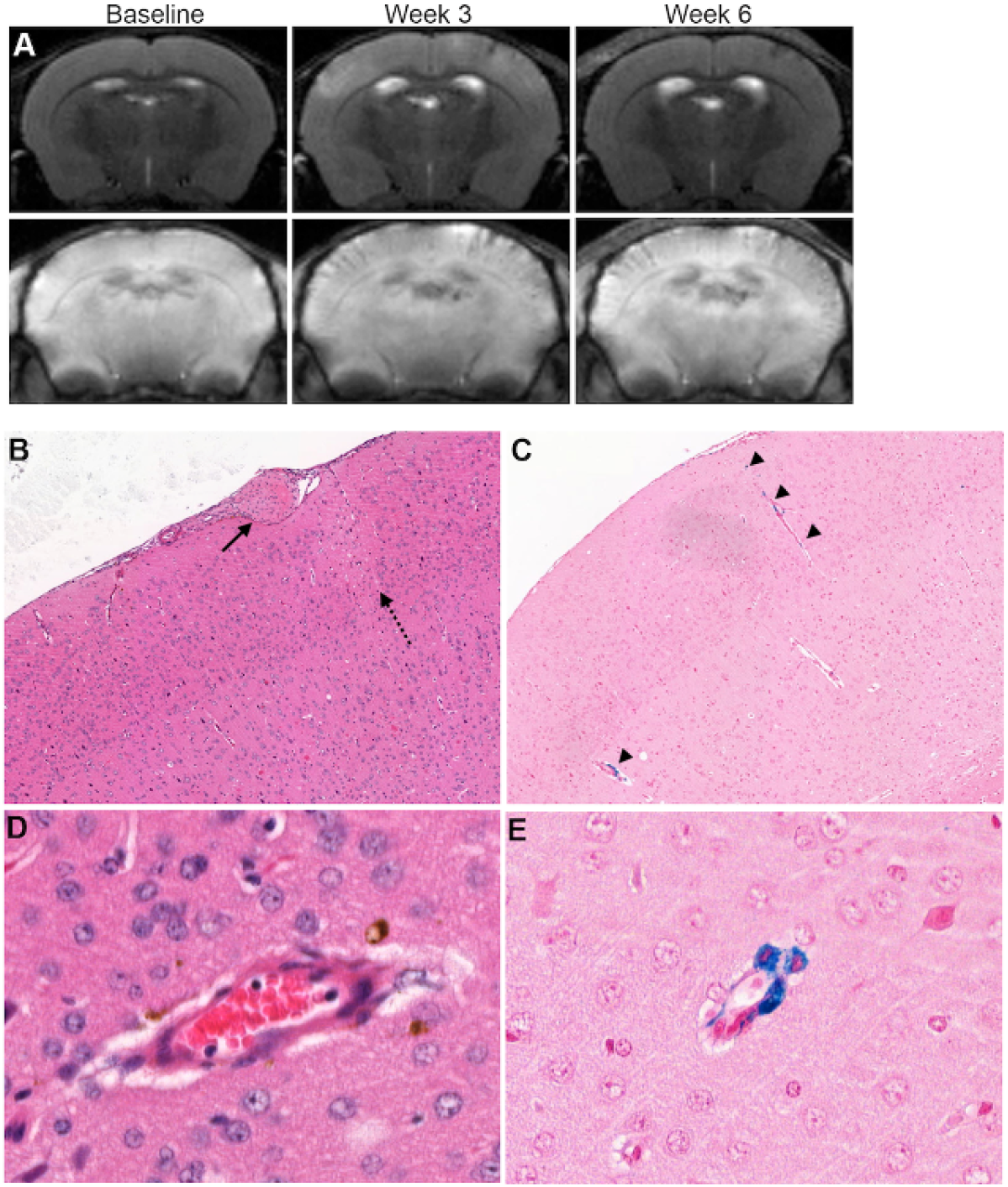
T2w-hyperintense, T2*-hypointense ARIA-like lesions observed in 5xFAD mice treated with 3D6 mIgG2a that correspond to perivascular microhemorrhages. **(A).** T2-weighted (top row) and T2*/GRE (bottom row) MRI scans (slice 16) from Animal 21/study 2, treated with weekly doses of 3D6 (10 mg/kg), showing: emergence of bilateral diffuse T2w-bright cortical lesions after dose 3 (week 3) along with T2*-dark cortical lesions that were linear and oriented perpendicular to the leptomeningeal surfaces, consistent with periarteriolar microhemorrhages. T2*-dark cortical lesions persisted after the 6^th^ dose (week 6) despite reduction of the diffuse T2w-bright cortical lesions. One T2w-bright left cortical lesion is also noted to persist at week 6. H&E-stained (**B, D**) and Perls iron-stained (**C, E**) sections from Animal 21. Vascular thrombosis (arrow), a rare finding in all studies, and acute microinfarct (dashed arrow) was observed in the right hemisphere (B, original magnification 20x). Perivascular hemosiderin was observed around several penetrating cortical arterioles (arrowheads) (C, original magnification 20x). High-power images of a penetrating arteriole with Perls iron-reactive perivascular hemosiderin (D, E; original magnifications 400x).

### ARIA-related vascular inflammation is associated with vascular leakage and microvascular lesions

To determine whether MVI is associated with compromised mural integrity and leakage of vascular luminal contents that underlie T2w-hyperintense MRI signals, spatial distributions of MVI and parenchymal albumin immunoreactivity (alb-ir) were compared in Study 2 mice. In a 3D6 mIgG2a-treated mouse with right-sided cortical ARIA-E-like lesions at week 8 just prior to necropsy, albumin IHC demonstrated abnormal diffuse parenchymal immunoreactivity in the right frontal and temporal cortices (**Fig. 8A, B**). In cortical regions with MVI, highlighted with Pu.1/Iba1 immunohistochemical detection of meningovascular macrophages, colocalization with parenchymal alb-ir was observed (**Fig. 8C, D**), whereas cortical regions lacking MVI (left temporal cortex) did not show abnormal parenchymal alb-ir (**Fig. 8E, F**). Accentuation of superficial/subarachnoid alb-ir and diminution of alb-ir in the deeper cortex, consistent with a meningovascular source of abnormal albumin brain permeation, were frequently observed (**Fig. 8G**). Perls iron histochemistry was employed (**Fig. 8H, I**) to highlight and enumerate perivascular hemosiderin deposits/foci which were frequently found in cortical alb-ir regions (**Fig. 8G I**). Cortical alb-ir, quantified as % Area of cortex immunoreactive for albumin, trended higher in the 3D6 mIgG2a treatment group of study 2 compared to baseline and control groups (**Fig. 8J**). In contrast, perivascular hemosiderin foci were observed only in the 3D6 mIgG2a treatment group (**Fig. 8K**). Even though there are temporal differences – alb-ir reflects transient and recent vascular injury and leakage, whereas Perls iron microhemorrhage may reflect vascular injury and leakage that can occur over the entire treatment period – we found a correlation between cortical alb-ir and cortical perivascular hemosiderin foci (Spearman r = 0.7230; p<0.0001).

**Figure 8.**
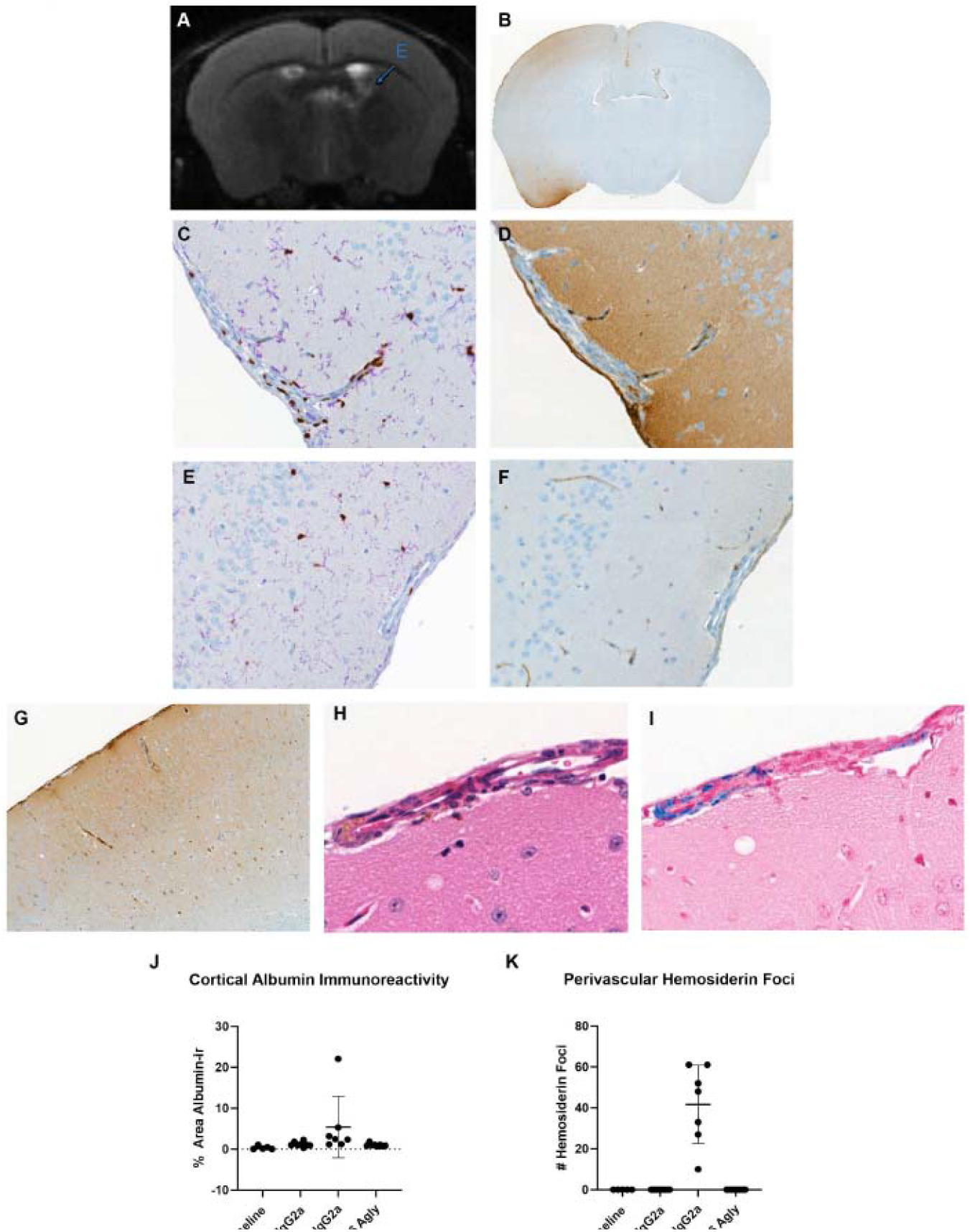
Abnormal vascular permeation and perivascular microhemorrhages in 5xFAD mice with ARIA-related MVI. **(A).** T2-weighted image showing superficial T2 bright signal involving the right frontal cortical convexity correlates with **(B)** parenchymal albumin immunoreactivity indicative of abnormal vascular permeation. Note that albumin immunoreactivity is most intense at the subarachnoid space and superficial cortex and wanes in the deep cortex, consistent with meningovascular permeation as the source. (**C)** Pu.1/Iba1 immunohistochemistry highlights right temporal MVI coinciding with **(D)** superficial cortical albumin permeation whereas, contralaterally, the lack of left temporal MVI (**E**) coincides with a lack of vascular albumin permeation (**F**). (**G)** Corresponding to the right middle-frontal region of T2-bright signal and vascular permeation by albumin, perivascular hemosiderin evident on H&E (**H**), and highlighted by Perls iron stain (**I**), is observed. (**J)**. Albumin immunoreactivity throughout cortical regions of interest trends higher in 3D6 IgG2a treated mice of study 2 compared to control IgG2a and 3D6-agly IgG1 treated mice. (**K)**. Leptomeningeal perivascular hemosiderin foci appeared only in 3D6 IgG2a treated mice with ARIA and were not found in untreated mice (Baseline), control IgG2a and 3D6-agly IgG1 treated mice.

### 3D6 induces vascular amyloid removal and is associated with vascular complement activation

We hypothesized that MVI represents antibody-dependent activation of perivascular macrophages following vascular Aβ engagement. Similar to previously published studies (*9, 11*), we observed that 3D6 treatment leads to vascular Aβ removal by macrophages (**Fig. 9A F**). Evidence of vascular Aβ engagement by 3D6 was further examined in meningeal arterioles lacking inflammation or showing low MVI. Near adjacent histologic sections were stained for Aβ 12F6A IHC (**Fig. 9G, I, K**) and mIgG IHC (**Fig. 9H, J, L**), showing that Control mIgG2a did not engage CAA (**Fig. 9G, H**), whereas 3D6 mIgG2a produce strong labeling of CAA (**Fig. 9I, J**), in the background of diffuse mIgG immunoreactivity due to vascular permeation associated with treatment-induced inflammation. While the binding to CAA was still observed with 3D6 mIgG1^agly^ (**Fig 9K**), the aglycosylated nature of the antibody and its reduced effector function is most likely responsible for the tamed inflammatory reaction. Notably, 3D6 mIgG1^agly^-treated mice lacked evidence of parenchymal mIgG immunoreactivity (**Fig 9L**) and ARIA-like lesions by MRI. Vascular profiles with inflammation (**Fig. 9M**) showed immunoreactivity for complement C1q (**Fig. 9N**) and C3 (**Fig. 9O**), suggesting that treatment-induced immune complex vasculitis could occur.

**Figure 9.**
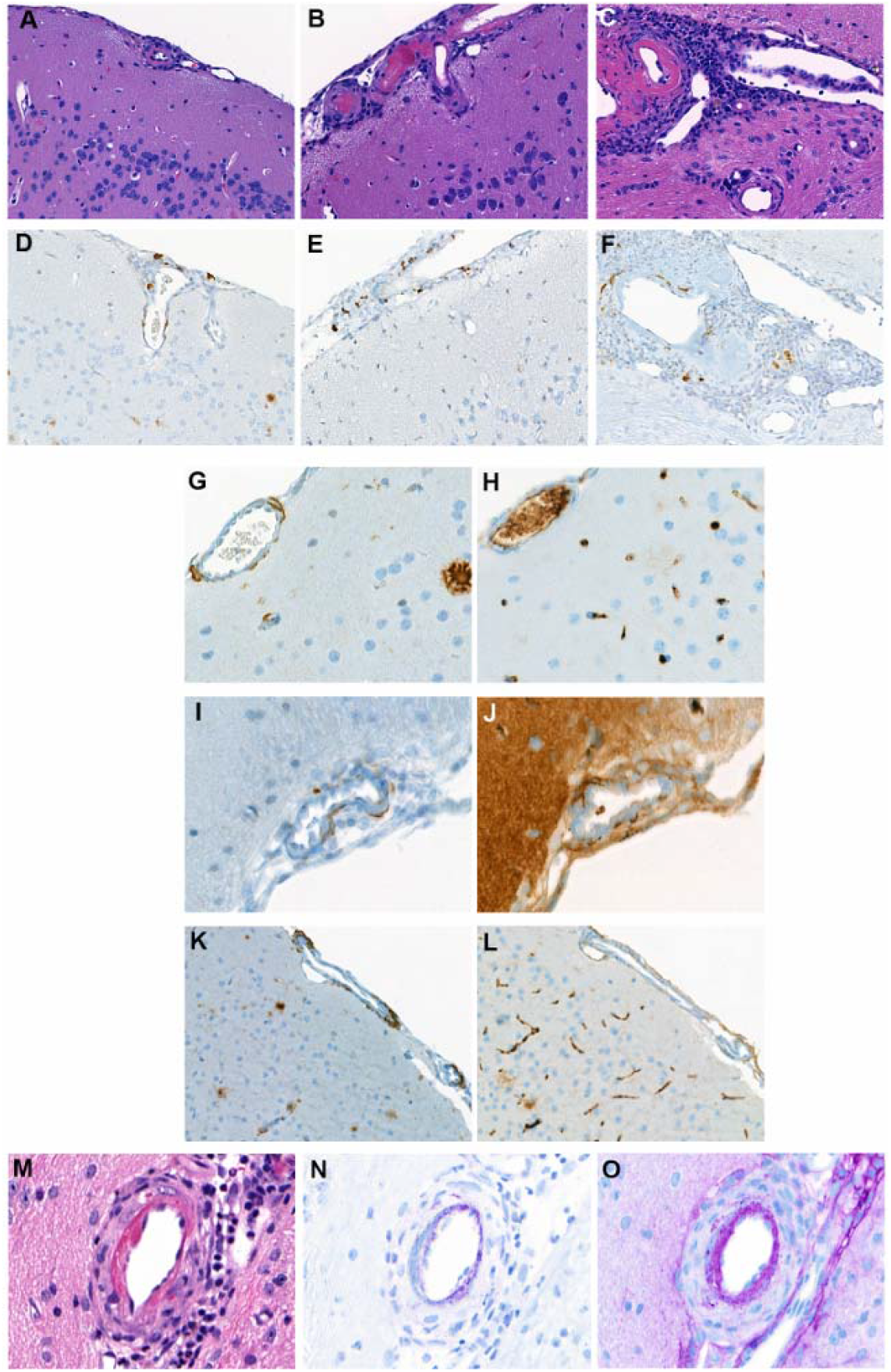
3D6 engages cerebrovascular Aβ (CAA) and induces MVI with complement cascade activation. A-F: Representative H&E sections with moderate **(A)** to severe MVI (**B-C**) and Aβ immunohistochemistry (12F6A; **D-F**) in near-adjacent sections demonstrating breakup, phagocytosis and clearance of vascular Aβ in macrophages / mononuclear infiltrates. G-L: In regions with minimal MVI, comparison of 12F6A immunohistochemistry (**G, I and K**) with murine IgG immunohistochemistry (**H, J and L**) in near-adjacent sections of arterioles. 12F6A IHC revealed amyloid angiopathy involving meningeal arterioles of 5xFAD mice that lack moderate to severe inflammation (**G, I and K**). Control mIgG2a shows absence of CAA engagement (**G, H**). Engagement of Grade 1 (adventitial) CAA by 3D6 mIgG2a (**I, J**) and 3D6 mIgG1^agly^ (**K, L**) are shown. **M-O:** Severe meningovascular inflammation with fibrinoid change (**M**; H&E) showed immunoreactivity for C1q (**N**) and C3 (**O**), consistent with activation of the classical complement cascade.

### ARIA-related vascular inflammation and leakage increase anti-Aβ antibody brain exposure, amyloid plaque engagement and clearance

The impairment of vascular integrity with 3D6 mIgG2a treatment raises the possibility that ARIA may lead to increased brain exposure of the administered anti-Aβ antibody. This possibility was investigated histologically by comparing the brain content of 3D6 mIgG2a which causes ARIA versus 3D6 mIgG1^agly^ which has the same molecular weight and binding properties as 3D6 mIgG2a, but does not induce ARIA. Both diffuse parenchymal and plaque-associated mIgG-immunoreactivity were higher in 3D6 mIgG2a treated mice with ARIA compared to 3D6 mIgG1^agly^ treated mice which did not have ARIA (**Fig. 10A J**). This even though Aβ plaque burden trended lower in the 3D6 mIgG2a treatment group compared to control mIgG2a or 3D6 mIgG1^agly^ treated mice (**Fig. 10K**). Moreover, there was a trend for an inverse linear correlation between Aβ plaque burden and cortical albumin immunoreactivity (**Fig. 10L**; Spearman r = - 0.7143, p=0.0881) and a strong inverse linear correlation between Aβ plaque burden and leptomeningeal microhemorrhages detected on Perls iron stain (**Fig. 10M**; Spearman r = -0.8214, p=0.0341) in 3D6 mIgG2a treated mice. This data supports the notion that when indicators of ARIA-associated vascular disruption were high, brain exposure and amyloid plaque targeting were increased and amyloid burden was ultimately reduced.

**Figure 10.**
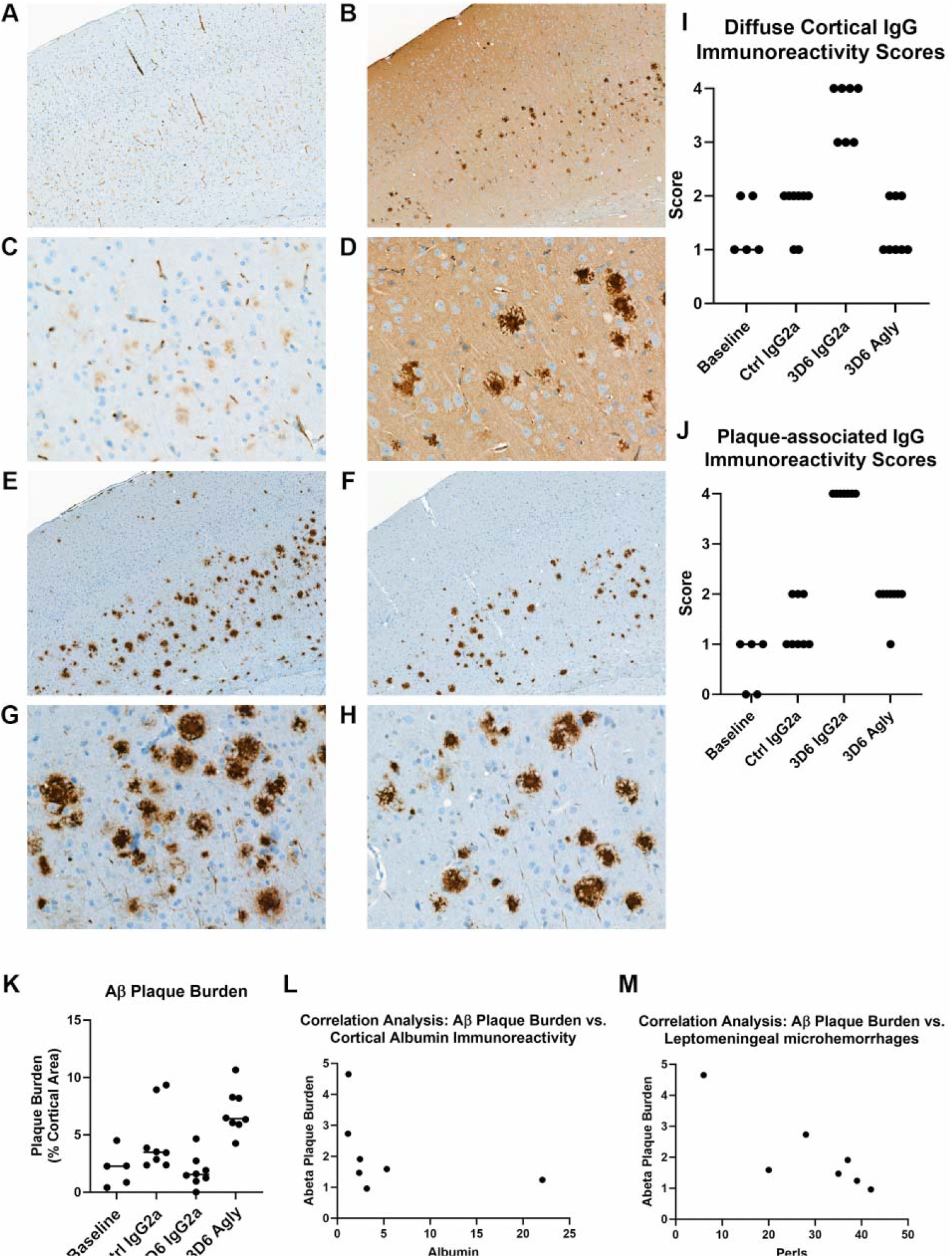
ARIA-related vascular inflammation and leakage augments brain and amyloid plaque exposure to 3D6. **(A-D)** Leakage of vascular content is evidenced by diffuse parenchymal murine IgG immunoreactivity and increased amyloid plaque-associated murine IgG immunoreactivity in sections from animal 11/study 2 with ARIA induced by 3D6 mIgG2a treatment (B, original magnification 50x; D, original magnification 400x), as compared to sections from animal 7/study 2 treated with 3D6 mIgG1^agly^ that did not show ARIA and did not exhibit parenchymal mIgG immunoreactivity (A, original magnification 50x; C, original magnification 400x). (**E-H).** representative images showing the Aβ plaque burdens (Aβ 12F6A IHC) in near-adjacent sections to those depicted in Panels A-D, from animal 7 without ARIA (E, G) and animal 11 with ARIA (F, H) (E, F; original magnification 50x; G, H; original magnification 400x). (**I-J)**. Semiquantitative murine IgG immunoreactivity scores indexing diffuse cortical IgG immunoreactivity (I) and plaque-associated IgG immunoreactivity (J) were elevated in 3D6 mIgG2a treated mice compared to control mIgG2a and 3D6 mIgG1^agly^ burdens in 3D6 IgG2a treated mice were lower compared to 3D6 mIgG1^agly^ treated mice. (**K)**. Aβ plaque treated mice (p=0.0020) and trended lower in 3D6 mIgG2a treated mice compared to control isotype mIgG2a treated mice (p=0.2003). (**L-M)**. A trend of an inverse linear correlation between Aβ plaque burdens and cortical albumin immunoreactivity (L, Spearman r = -0.7143, p=0.0881) and a strong inverse linear correlation between Aβ plaque burdens and leptomeningeal microhemorrhages detected on Perls iron stain (M, Spearman r = - 0.8214, p=0.0341) was observed in 3D6 mIgG2a treated mice.

### Incidence and severity of ARIA-like lesions seen in 5XFAD mice with murine versions of bapineuzumab, aducanumab and gantenerumab parallel results seen in clinical trials

Considering the higher incidence and severity of ARIA observed in clinical trials with bapineuzumab, the characterization of the model was carried out by treating 5xFAD mice with the murine precursor of bapineuzumab 3D6. Additional studies were then conducted with gantenerumab and aducanumab in order to validate the model. While the imaging characteristics of ARIA-like lesions were similar for all 3 antibodies, the incidence of the lesions varied between treatment. ARIA-like lesions were observed in only 25% of 5xFAD mice treated with ^ch^gantenerumab mIgG2a at 50 mg/kg with delayed onset compared to 3D6 and was not observed at 5, 10 or 30 mg/kg doses (studies 4-6, Table 1; Fig. 5). In limited testing at 50 mg/kg (n=18 total), ARIA-like lesions were identified in approximately 50% of 5xFAD mice treated with aducanumab hIgG1 or murine chimeric aducanumab mIgG2a (study 7, Table 1; Fig. 5). Treatment with an aglycosylated version of either 3D6 or gantenerumab failed to induce any ARIA-like lesions, supporting a critical role for effector-function in the pathogenic mechanisms for ARIA. The differential incidence of ARIA-like lesions observed with the three different antibodies tested in the current mouse model is in line with clinical data of the corresponding anti-Aβ molecules and provides further support for the model translatability.

The model was leveraged to assess the potential risk for ARIA of a newly developed TfR-based anti-Aβ BBB-shuttle, demonstrating a low rate of ARIA with ATV:Aβ as compared to its non-shuttled equivalent 3D6 (*20*). Results from these preclinical studies are in agreement with the clinical data described with the TfR-based BBB-shuttle version of gantenerumab, trontinemab, showing an incidence of ARIA-E and ARIA-H lower than 5% after 28 weeks of treatment (*22*). Concordance of ATV:Aβ preclinical data with trontinemab clinical data further supports the validity of the current model to assess the next generation of Aβ immunotherapeutics. Histograms of ARIA prevalences and MVI severity grades among 5xFAD mice treated with 3D6, gantenerumab and aducanumab (**Fig. 5**) confirms a close association between MVI, in particular moderate to severe MVI, and ARIA. Notably, several animals in the gantenerumab and aducanumab groups demonstrated minimal to mild, and occasionally focally moderate, MVI without ARIA detection.

## DISCUSSION

Clinical trials with monoclonal antibodies targeting different forms of Aβ aggregates, such as aducanumab, lecanemab and donanemab, have shown reduction in brain amyloid burden and a slowing of disease progression in patients with early symptomatic AD, leading to their approval by certain regulatory authorities (*23*). This represents one of the major advances over the last decade in the quest for developing effective drugs for the treatment of Alzheimer’s Disease. However, their administration can lead to occurrence of Amyloid-Related Imaging Abnormalities (ARIA) in some patients, and ARIA still remains the main adverse event occurring with the anti-Aβ mAbs class of drugs.

Originally described as vasogenic edema in patients treated with bapineuzumab (*24*), these adverse events were formally identified and termed ARIA by the 2010 Research Roundtable Workgroup (*25*). Since then, ARIAs have been described with most of the anti-Aβ mAbs, and some of their key features have been further refined based on data from multiple clinical trials (*2, 26*). ARIA-E is detected as hyperintensity on T2w or FLAIR MRI sequences, and would correspond to leakage of proteinaceous fluids in the brain parenchyma as vasogenic edema or sulcal effusion in the leptomeningeal space. ARIA-H is often detected as hypointense signal on a T2*-GRE MRI sequence, and would be associated with leakage of blood degradation products in the parenchymal space as microhemorrhage or in the subarachnoid space as cortical superficial siderosis. While few spontaneous cases of ARIA (i.e., in the absence of anti-Aβ immunotherapy) have been reported, most ARIA cases are considered to be treatment-related adverse events, they occur early in the course of the treatment and often resolves upon cessation of the treatment. The majority of ARIA cases are clinically asymptomatic and detected through scheduled MRI monitoring. Higher dose levels of the drug, apolipoprotein E4 haplotype and presence of cerebral amyloid angiopathy (CAA) represent the major known risk factors for ARIA. As of today, many questions remain unanswered about the pathogenesis and clinical consequences of ARIA, and this is mostly due to the absence of predictive biomarkers, the absence of clear correlation between imaging and clinical severity, and the scarcity of correlative data between MRI findings and pathological observations, other than few rare but severe and fatal cases (*3-5*). While recent publications reported neuropathological examination of brains from subjects treated with anti-Aβ immunotherapy, only a couple of cases effectively presented with ARIA during the course of treatment, and autopsy for these cases was performed 32-38 months after the active ARIA events (*6, 26, 27*). In addition, preclinical research has often used the occurrence of microhemorrhages in amyloid-bearing mice treated with therapeutic candidates as the sole correlate of ARIA, hence possibly ignoring lesions such as edema that are often transient and do not always evolve into microhemorrhage.

Therefore, the development of a relevant ARIA mouse model recapitulating some of the ARIA features observed in human subjects treated with Aβ immunotherapy would be useful to i) investigate and identify ARIA pathogenic mechanisms; ii) explore means to mitigate ARIA; iii) reliably assess and derisk new therapeutic compounds under development. The mouse model described in the current manuscript utilizes clinically relevant MRI sequences to monitor “ARIA-like” events in amyloid plaque-bearing transgenic mice chronically treated with Aβ immunotherapy. Correlation of the imaging findings with histological assessments is used to identify molecular changes underlying ARIA and establish potential pathogenic mechanisms.

Our studies demonstrate that treatment with the murine parent of the first clinically tested anti-Aβ mAb bapineuzumab readily induced ARIA-like lesions in 5xFAD mice, most of which showed characteristics that resemble human ARIAs. T2 hyperintense lesions in these mice histologically corresponded to abnormal brain parenchymal albumin immunoreactivity and augmented brain parenchymal IgG immunoreactivity consistent with compromised vascular integrity and leakage of blood components (vasogenic edema). T2* hypointense lesions resembled ARIA-H lesions in patients, often occurring in regions previously involved by T2 hyperintensities. Histologically, perivascular T2* hypointense MRI lesions corresponded to Perls iron positive hemosiderin remnants of microhemorrhages around meningeal and penetrating cortical arterioles. The similarities between the ARIA-like lesions we observed in 5xFAD mice and human ARIA lesions suggest common pathologic mechanisms that might be discovered through the study of murine ARIA-like lesions in lieu of the limited availability of human tissue except from the most severe, fatal ARIA cases.

Our data demonstrate a strong association between the histopathologic finding of meningovascular inflammation (MVI) and ARIA-like lesions detected by MRI, suggesting that MVI is a key pathogenic driver of ARIA-like lesions in 5xFAD mice. Moreover, they support the hypothesis that systemically administered Aβ antibodies induce ARIA by engaging vascular Aβ deposits and recruiting an inflammatory response comprised predominantly of activated macrophages, a response that is absent with effectorless Aβ antibodies (*28-30*). Vascular inflammation compromises vascular mural integrity leading to vascular permeation and extravasation of plasma albumin, which is accentuated in a superficial distribution at the site of meningovascular inflammation, and erythrocytes. Recent preclinical studies by several groups have implicated parenchymal border macrophages in Aβ antibody-induced vascular injury (*12, 14, 31*) and further implicated macrophage interactions with APOE4 (*31, 32*), a genetic risk factor for CAA and ARIA. Notably, Trem2-positive macrophages which were implicated by Taylor and colleagues in microhemorrhages in PDAPP mice treated with 3D6 (*14*) also may play an important role in human ARIA as suggested by ARIA induction in AD patients treated with TREM2 activating antibodies (*33*). Preclinically, complement activation has also received attention as a potential effector of inflammation and vascular damage (*34*), though its role in early vascular inflammatory lesions needs to be studied further to justify a prevention strategy based on complement cascade inhibition. Human neuropathologic studies of both rare fatal cases of ARIA (*3-5*), spontaneous CAA-ri (*35*) and spontaneous CAA-associated microbleeds (*36-39*) also implicate neuroinflammation in response to vascular amyloid resulting in vessel remodeling with Aβ removal and hemorrhage (*40*). Moreover, a prevailing hypothesis is that ARIA represents iatrogenic CAA-ri which might be associated with elevated CSF levels of Aβ autoantibodies. Our study fills a gap by establishing a strong link between these pathologies and MRI-evidence ARIA-like lesions in a translational model of ARIA.

From a translational viewpoint, our data suggest that induction of ARIA-like lesions in 5xFAD mice with 3D6 should be used for the preclinical evaluation of next generation Aβ- targeting therapies with ARIA-mitigation strategies (*20*). Notably, the histologic assays employed in this study showed high concordance with MRI findings. Our blinded meningovascular inflammation scoring method leveraging H&E-stained sections showed excellent concordance with ARIA-like events on MRI. Perls iron has been employed by several groups in murine Aβ models with the assumption that microhemorrhages inform on ARIA-like events (*9, 12, 14, 41, 42*). In our study, Perls iron histochemistry showed the best sensitivity and specificity for ARIA-like lesions, likely due to the persistence of tissue hemosiderin and wide sampling of cortical meninges. Our method of blinded scoring has an advantage over color deconvolution-based image analysis due to the known noisy artifacts that can occur with Perls iron histochemistry particularly at the leptomeningeal tissue edges, however, the latter can be employed when artifacts are sparse. Albumin immunohistochemistry showed good specificity but low sensitivity in this study perhaps due to the evolution/resolution of vascular permeation. Taken together, our data provide support that a suite of assays (H&E, albumin and Perls iron) can provide surrogate markers for ARIA-like events in 5xFAD mice when MRI is not feasible for preclinical studies.

In contrast to biweekly or monthly dosing schedules utilized in AD patients, we employed a weekly dosing protocol that resulted in ubiquitous induction of ARIA-like lesions in 3D6 mIgG2a-treated 5xFAD mice within 3-8 weeks. While it is potentially advantageous to have robust ARIA-like lesions for preclinical testing of therapeutics that may mitigate ARIA, the frequency and severity of the lesions we observed with weekly dosing of 3D6 exceed those seen in the clinic and thus generalizability of the findings to human ARIA may have pitfalls. Notably, severe meningovascular inflammatory lesions were common in our 5xFAD mice and occasional parenchymal lesions, predominantly microinfarcts that are indicative of compromised blood flow secondary to meningovascular inflammation, were observed. Microinfarcts have thus far not been observed or attributed to anti-Aβ treatment in humans (no restricted diffusion; no evidence in few autopsies AN1792). Examination of bapineuzumab-induced ARIA lesions were reported to show no restricted diffusion (ischemic features) (*43*). However, small regions of restricted diffusion suggestive of the possibility of small ischemic infarcts have recently been reported in a patient treated with lecanemab (*44*), suggesting that ischemic events can occur with amyloid removal therapy but are rare. The meningovascular inflammatory responses and parenchymal changes we observed in 5xFAD mice arguably show resemblance to severe meningovascular inflammatory findings in rare fatal ARIA cases (*3-5*) but the generalizability to most ARIA events in AD patients is unclear and will likely remain so due to lack of tissue accessibility during active ARIA.

We also observed evidence of higher brain and cortical Aβ plaque exposure to 3D6 IgG2a compared to 3D6 IgG1^agly^ due to meningovascular inflammation and permeation. In support of a significant effect of this higher exposure on Aβ plaque clearance, we observed a strong inverse correlation between the burden of microhemorrhages (revealed by Perls iron stain) and Aβ plaque burden among 3D6 IgG2a-treated 5xFAD mice. A trend towards a strong inverse correlation between the cortical albumin immunoreactivity and Aβ plaque burden was also observed. Cortical Aβ plaque burden was reduced with only 8 weeks of 3D6 IgG2a treatment. These data support augmented brain exposure to antibody with ARIA and raise the consideration that while ARIA is considered undesirable, its potential to expose the brain to more rapid effects on amyloid clearance may have a confounding factor for amyloid clearance and efficacy.

Our data also clearly show that anti-Aβ effector function is a major determinant of ARIA risk in 5xFAD mice, supporting the possibility that effector function tuning to optimize reactivity with Aβ plaques and minimize reactivity with vascular Aβ could mitigate ARIA risk. Clinical data obtained with AAB-003, a modified version of bapineuzumab with reduced Fc-receptor-mediated effector function, provides further validation of these observations (*45*). Moreover, lower rates of ARIA with gantenerumab and aducanumab compared to 3D6 in 5xFAD mice implicate important contributions of antibody variable region to ARIA risk, though ARIA rates for gantenerumab and aducanumab in the clinic were still high. A recent study implicated antibody binding affinity to CAA as an important determinant of ARIA risk in patients (*46*). Further understanding of these factors may aid in the design of novel ARIA mitigation strategies.

The 5xFAD model develops abundant Aβ plaque pathology and shows Type II CAA at 3-4 months of age (*19, 21*), however it is not considered a robust model of CAA. Nonetheless, we found that ARIA-like lesions and MVI can readily be induced in 5xFAD mice ranging from 7-15 months of age. Our work did not examine the potential contribution of paravascular Aβ clearance which has been hypothesized to underlie a shift of Aβ from plaques to CAA following Aβ immunotherapies in patients (*47, 48*). Moreover, we did not evaluate a potential contribution of neuroinflammation-induced blood brain barrier permeation to the pathogenesis of ARIA-like lesions. Our studies were primarily designed to establish a murine model that could be leveraged to evaluate therapeutics that mitigate ARIA and were not optimally designed to dissect underlying mechanisms which could benefit from earlier necropsies before, during and immediately after the onset of ARIA-like lesions.

In summary, our radiologic-pathologic correlation study leveraging human-relevant MRI sequences demonstrates that 5xFAD mice treated with anti-Aβ immunotherapy develop ARIA-like lesions that bear considerable similarity to human ARIA lesions though may represent the more severe side of the spectrum. These lesions correspond to neuropathologic changes including meningovascular inflammation and vascular permeation, and they expose the brain and Aβ plaques to higher levels of anti-Aβ than seen in the absence of ARIA and may increase rate of Aβ plaque removal. This model has been useful for the preclinical evaluation of next generation Aβ targeted therapies with ARIA mitigation strategies. Although the limited access to small animal MRI scanners could represent a significant burden to a broad use of the ARIA murine model, one could envision additional studies to fully exploit the benefits of this model to further understand human ARIA mechanisms. For example the choice of the mouse model is critical as it needs to finely mimics human amyloid pathology, including parenchymal plaques and CAA, or the dependance towards ApoE4-related phenotype. This type of preclinical model is also appropriate to investigate the potential role of different cell types, neurimmune interactions, or vascular factors involved in the pathogenesis of ARIA. And finally, this model provides a significant advantage in terms of possible adjustment of the treatment paradigm or the timeline for MRI acquisition and corresponding histopathology.

## MATERIALS AND METHODS

### Generation of anti-Aβ Antibodies

Published amino acid sequences of the variable domains of 3D6 (the murine parent of bapineuzumab) (*49*) and gantenerumab (*50*) were used to generate different variants of each antibodies. These included: (i) a murine IgG2a version of 3D6 (3D6 mIgG2a); (ii) a murine aglycosylated IgG1 version of 3D6 (3D6 mIgG1^agly^); (iii) a murine chimeric IgG1 version of 3D6 (^ch^3D6 hIgG1); (iv) a murine chimeric IgG2a version of gantenerumab (Gante mIgG2a); and (v) a murine chimeric aglycosylated IgG1 version of gantenerumab (Gante mIgG1^agly^). These antibodies (3D6 mIgG2a, 3D6 mIgG1^agly, ch^3D6 hIgG1, and Gante mIgG1^agly^) contain the variable heavy (V_H_) and variable light (V_L_) domains of the specific antibodies and murine IgG2a/kappa, murine IgG1 with N297A mutation (Agly variants) and human IgG1 constant heavy chain and constant light chain domains for the human variants. The murine chimeric IgG2a/kappa version of aducanumab (^ch^Adu) was generated as described (*51*) and tested alongside unmodified human aducanumab. All antibodies were expressed in CHO cells and purified by protein A affinity followed by ion-exchange chromatography.

### Animal care and in vivo procedures

5xFAD (strain B6SJL-Tg(APPSwFlLon,PSEN1*M146L*L286V)6799Vas/Mmjax) male transgenic mice were purchased from The Jackson Laboratory at 2 months of age and were housed in the vivarium facility at Biogen until they reach the required age. Genotype was confirmed by analyzing DNA extracted from tail clippings (Transnetyx, Cordova, TN). Mice were housed with a 12-hour light/dark cycle, and food and water were provided ad libitum. Aβ antibodies were administered weekly via intraperitoneal injections using a 30G-gauge needle, for 5-12 weeks. **Table 1** summarizes studies protocol for 7 Study cohorts (n=153 5xFAD and WT mice total) analyzed in this current report. All experimental procedures were approved by Biogen Institutional Animal Care and Use Committee (IACUC) and adhered to all applicable guidelines and regulations (Cambridge Ordinance 1086, PHS Policy, AWA/AWAR, and the Guide) and the latest IACUC Guidance for the Humane Euthanasia of Laboratory Animals guidelines.

### MRI protocols and analysis

Mice were anesthetized with isoflurane in medical oxygen (2% induction, 1-2% maintenance). Scans were acquired on a 9.4T preclinical MRI (Bruker BioSpec 94/20, Billerica, MA, USA) equipped with Paravision (version6.0.1). A mouse brain receive-only phased array surface coil was positioned over the head for signal reception, and an 86 mm actively detuned transmit-only volume coil was used for transmission. Animals were secured in the head holder using a bite bar and ear bar. Respiration was maintained between 70-160 breaths per minute.

Body temperature was maintained at 37 □ ± □ 0.3 °C using an MR-compatible warm air heating system (Small Animal Instruments, Inc. Stony Brook, NY, USA). Temperature was monitored via a rectal probe. Baseline scans were acquired to document the absence of pre-treatment abnormalities and thereafter, scans were acquired throughout the duration of studies within 1-3 days after weekly antibody administrations. T2-weighted (T2w) rapid acquisition relaxation enhance (RARE) image was acquired (slice thickness= 0.5mm, repetition time [TR]= 4100ms, echo time [TE]=60 ms, averages=10, repetitions=1, echo spacing=7.5ms, rare factor=16, flip angle=90° FOV=16x16mm, matrix= 128x128, slices=30) followed by 2D multislice Multi-Gradient Echo (MGE) image (slice thickness=15mm, TR=70ms, TE=3ms, averages=1, repetitions=1, echo images=12, echo spacing=4.5ms, flip angle=30°, FOV=16x16x15mm, matrix= 128x128, slices=30). For certain studies diffusion-weighted imaging (DWI) was performed on animals that exhibited T2-hyperintensities (slice thickness=0.5mm, TR=3200m, TE=21.5ms, averages=1, repetitions=1, segments=2, flip angle=90°, FOV= 16x16mm, matrix=128x60, slices=30, 3 orthogonal diffusion directions with B-values b=0, 500, 1000 s/mm^2^). T2w, T2* and DWI weekly scans were reviewed for presence of ARIA-like lesions, i.e., any new MRI event not present on baseline scan. Descriptions of lesions include information about imaging features, location and temporal evolution. The presence of ARIA-like lesions was determined by investigators (PCTC and JA) blind to the treatment animals received and these results were reviewed by two additional independent investigators (DB and TB).

### Histopathology assays and analysis

Animals were deeply anesthetized with 1.9L CO_2_/min and transcardially perfused with heparinized saline followed by 4% paraformaldehyde. Brains were carefully removed to preserve the leptomeninges and were postfixed in 10% neutral-buffered formalin for 48-72 hours. Fixed brains were marked with red or green ink on the right hemisphere and then coronally sectioned at 2 mm intervals with the first cut at the optic chiasm. Six coronal slices were processed and embedded in paraffin blocks for sectioning in the rostral-to-caudal direction. Blocks were gently faced and sectioned at 5 μm thickness to produce 10 consecutive unstained slides. One slide was stained with hematoxylin and eosin (H&E) and one slide was stained with the Perls iron method (Prussian Blue Iron Kit; Polysciences Inc.; Cat # 24199). The following primary antibodies and assay conditions were employed in immunohistologic assays performed on Ventana Discovery Ultra autostainers: Aβ (clone 12F6A, epitope 3-7; Biogen, 0.25 μg/mL; diluted in antibody diluent with casein, CC1 antigen retrieval for 64 minutes, a blocking step in Protein Block (Dako Cat# X0909) prior to antibody incubation and a linking Rabbit anti-human IgG antibody (1 μg/mL, Jackson Immuno Cat # 309-005-003) after 12F6A incubation; rabbit anti-albumin (Abcam Cat # ab192603; 0.5 μg/mL; CC1 antigen retrieval for 64 minutes); murine IgG (anti-mouse OmniMAP HRP; Roche Cat # 760-4310; CC1 antigen retrieval for 64 minutes); rabbit anti-C1q (Abcam Cat # ab182451; 5 μg/mL; Protease antigen retrieval for 8 minutes); rabbit anti-C3 (Abcam Cat #ab200999; 0.5 μg/mL; CC1 antigen retrieval for 64 minutes). A duplex immunohistochemistry assay utilizing rabbit anti-Pu.1 (clone 9G7; Cell Signaling Cat # 2258; 1.5 μg/mL) and rabbit anti-Iba 1 (Wako Cat # 019-19741; 0.125 μg/mL) was employed to visualize microglia. A CC1 antigen retrieval for 64 minutes was used to unmask the epitopes. The Pu.1 antibody was applied first in the dual sequence, using an HQ-HRP amplification secondary antibody to detect and DAB to visualize. A neutralization step was performed before the second antibody, Iba-1, was applied using an anti-Rb-HRP to detect and Discovery purple to visualize. Slides were coverslipped on a Tissue-Tak Glas automated glass coverslipper (Sakura Finetek). Whole slide images were acquired on a Pannoramic P250 slide scanner with a 20x objective.

All histologic analyses were performed by an investigator (EDP) blind to the treatment `animals received` and the MRI results. H&E slides were examined for meningovascular inflammation that was graded according to the following system: Grade 0 (none) – no obvious meningovascular inflammation; Grade 1 (minimal) – sparse perivascular mononuclear cells without obvious phagocytic morphology – when present, a focal or multifocal finding; Grade 2 (mild) – subtle vascular macrophage infiltrates requiring high power examination to visualize; Grade 3 (moderate) – multiple layers of vascular macrophages that were evident at low-to-medium power; Grade 4 (severe) – any vascular inflammatory infiltrate with clear fibrinoid change. Grades were assigned to all cortical arteriolar profiles in the 6 brain sections that showed inflammation. Meningovascular inflammation data were: (1) typically summarized on a per animal basis as the predominant/maximal meningovascular inflammation grade; or (2) in Cohort 1, summated over all 6 brain sections for a summated meningovascular inflammation score. H&E-stained slides were also examined for parenchymal lesions, including microinfarcts and hemorrhages.

Perls iron slides were examined for microhemorrhages. Leptomeningeal vascular profiles demonstrating 3 or more clear Perls-reactive hemosiderin foci and that were confirmed on H&E to contain hemosiderin pigment were counted (leptomeningeal hemosiderin foci). Perivascular hemosiderin foci around penetrating cortical arterioles were counted separately (penetrating perivascular hemosiderin). Finally, microinfarcts that showed any hemosiderin staining were designated as hemosiderin-positive microinfarcts and counted.

Murine IgG immunohistochemistry assays were analyzed using a semiquantitative visual scoring system for diffuse cortical immunoreactivity and for Aβ-plaque associated immunoreactivity. For diffuse cortical immunoreactivity, a score of 1 was assigned if the cortex appeared blue at low power; a score of 2 was assigned if the cortex was predominantly light brown at low power; a score of 3 was assigned if the cortex was at least focally dark brown at low power; a score of 4 was assigned if the cortex was predominantly dark brown at low power. For Aβ-plaque associated immunoreactivity, a score of 1 was assigned if, at low power, faint minimal plaque labeling was evident; a score of 2 was assigned if plaque labeling was increased; a score of 3 was assigned if, at low power, high cortical plaque immunoreactivity was uniformly seen; a score of 4 was assigned if, at low power, high cortical plaque immunoreactivity was uniformly seen.

Tissue section immunoreactivity was analyzed with custom-designed, threshold-based segmentation protocols using median filters and respective color deconvolution filters in Visiopharm Image Analysis Software (version 2019.12.0.6842 and subsequent updates). Cortical albumin immunoreactivity was calculated as the summed areas of cortex immunoreactive for albumin divided by the composite cortical area.

### Data Analysis

Histopathology assay data were plotted and analyzed using GraphPad Prism (v. 10.1.2). Multiple group comparisons of normally distributed data were conducted with ANOVAs. Mixed model logistical regression was employed to model the contributions of meningovascular inflammation and study number to ARIA risk in all mice. Time to ARIA onset in all mice was plotted using the survival analysis module of GraphPad Prism (v10.1.2). MVI inflammation sensitivity and specificity analysis for ARIA were conducted in GraphPad Prism (v10.1.2).

A logistic mixed effects model was used to quantify the risk of ARIA-E as a function of MVI grade, with MVI grade (categorical with levels of 0-1, 2, 3, and 4) the only fixed effect, and between-experiment heterogeneity accounted for using a random intercept for experiment ID. Model-based probabilities of ARIA-E for each MVI level were computed with 95% confidence intervals (CIs). Odds ratios with 95% CIs were computed for all pairwise combinations of MVI levels; the Tukey HSD method was used to adjust odds ratio CIs for multiple comparisons. Modeling was conducted in R, with the lme4 used to fit the model, and the emmeans package used to compute probabilities and effects of interest.

## List of Supplementary Materials

Fig. S1 to S4 for multiple supplementary figures

## Funding

Research was funded by Biogen

## Author contributions

Conceptualization: EDP, DB, TB

Methodology: EDP, JA, PCTC, MLB, ACSA, JS, SH, JS, RB, FQ, DB, TB

Investigation: SBB, DLA, MPW, WCB

Data analysis: EDP, JA, PCTC, MLB, ACSA, SH, RB, DB, TB

Supervision: EDP, TB

Writing – original draft: EDP, JA, MLB, TB

Writing – review & editing: EDP, JA, PCTC, MLB, FQ, PW, DMW, DB, TB

## Competing interests

All authors either are current employees and shareholders of Biogen or were employees and shareholders of Biogen when these studies were conducted.

