## Supplemental Figures for "An in vivo model of Amyloid-Related Imaging Abnormalities (ARIA) predicts ARIA incidence of anti-Aβ antibodies seen in clinical trials"

**Affiliations:**

Biogen

Cambridge, MA, USA

*Corresponding authors:


### Supplemental Material

**Figure S1. Survival analysis – time to first ARIA event in all mice of 7 experimental cohorts.**

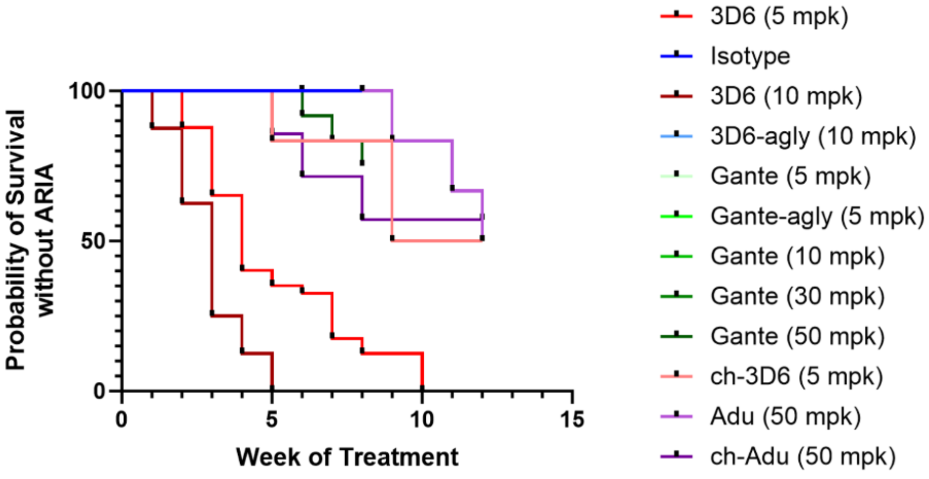


**Figure S2. Acute MVI with neutrophils and fibrinoid change was rarely observed when onset of ARIA was late in the dosing schedule and close to the time of necropsy.**

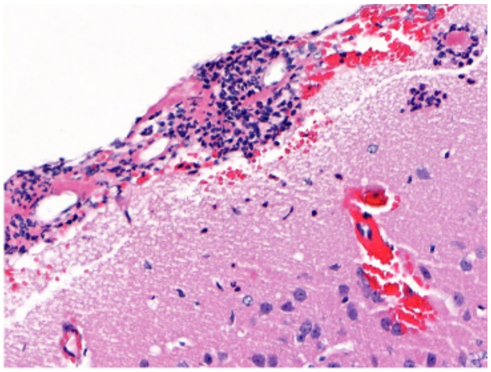

**Figure S3. Microinfarct counts in 5xFAD mice treated with anti-Aβ antibodies.**

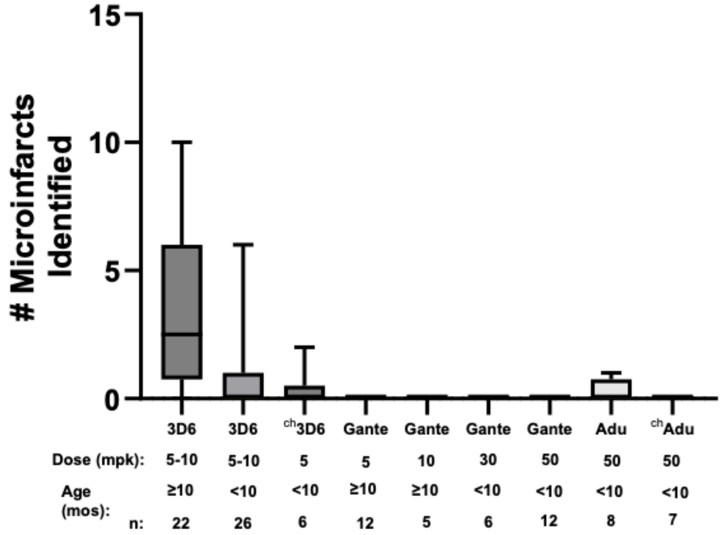


**Figure S4. H&E images from the largest meningovascular microhemorrhage observed in the studies. High power image in B shows acute neuronal necrosis and reactive gliosis, demonstrating this was a premortem hemorrhage.**

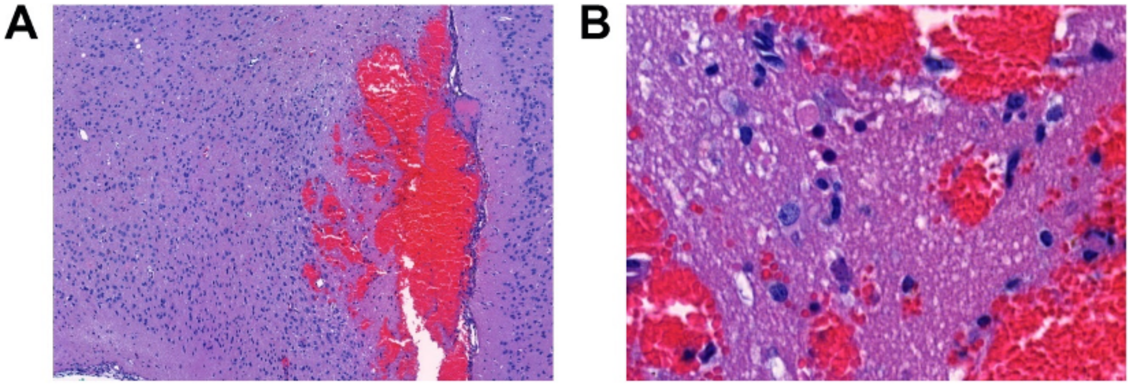
